# Phosphoinositide lipids have bidirectional and spatially distinct roles in filopodial dynamics

**DOI:** 10.1101/2025.10.13.681625

**Authors:** Julia Mason, Thomas C. A. Blake, Roshan Ravishankar, Oscar E. M. Despard, Gaudenz Danuser, Jennifer L. Gallop

## Abstract

Neural circuits are assembled by filopodia, dynamic actin-rich protrusions that connect environment sensing with structural changes. Using rapid timelapse imaging and acute pharmacological perturbation in *Xenopus laevis* retinal ganglion cells, we reveal that phosphoinositide lipid conversions impose spatially distinct control over filopodial dynamics. The class I phosphoinositide 3-kinase (PI3K) inhibitor alpelisib and Oculocerebrorenal syndrome of Lowe protein/Inositol polyphosphate 5-phosphatase inhibitor YU142670 markedly reduced filopodial tip motility. Class II PI3K inhibitor PITCOIN4 and YU142670 curtailed base dynamics, indicating location-specific lipid requirements. Despite no detectable change in PI(4,5)P₂, both alpelisib and YU142670 depleted PI(3,4)P₂ at filopodial tips. Their co-application was non-additive, with partial alleviation of YU142670-induced stalling by alpelisib, consistent with networked control of lipid conversion and actin remodelling. Live imaging with a TAPP1-3xcPH probe combined with quantitative cross-correlation and Granger causality analysis showed that PI(3,4)P₂ at filopodial tips both generates tip extension and responds to forward filopodial movement. Disrupting actin polymerisation rapidly erased tip PI(3,4)P₂ prior to filopodial stalling. These findings define a vulnerable node with implications for neurodevelopmental miswiring.

## Introduction

Filopodia are central to the assembly of neuronal circuits, acting as dynamic sensors and initiators of connectivity (Wit & Hiesinger 2022; Blake & Gallop 2023). These thin, actin-based protrusions extend from neuronal membranes to explore the extracellular environment, mediate initial contacts, and guide axonal growth cones toward their targets (Chien et al. 1993). In early development, dendritic filopodia are abundant and highly motile, gradually transforming into dendritic spines in response to synaptic activity (Ziv & Smith 1996). In axons, their regulated formation and stabilisation are essential for terminal arborisation and synapse formation (Dwivedy et al. 2007). To understand how neuronal circuits emerge, we must dissect how signals from adhesion, guidance cues, mechanical forces, synaptic activity, and ion transients converge to drive remodelling of the actin cytoskeleton (Thompson et al. 2019; Jain et al. 2024; Lew et al. 2025; Liu et al. 2025).

Retinal ganglion cell neurons in *Xenopus laevis* provide a powerful system for dissecting the cellular logic of axonal connectivity and miswiring as their stereotyped projection from retina to tectum enables precise analysis of pathfinding (Mann et al. 2004; Santos et al. 2020). Their large growth cones and long filopodia offer a rare opportunity to image filopodial dynamics in real time, despite the inherent challenges posed by their thin dimensions and rapid movement (Urbančič et al. 2017; Santos et al. 2020). These cells reflect general principles of circuit assembly that, when disrupted, can contribute to cognitive and behavioural symptoms in neurodevelopmental disorders (Bagni & Zukin 2019).

Phosphoinositide lipids have emerged as key intermediaries in the conversion of extracellular signals into cytoskeletal rearrangements (Janmey et al. 2018; Hou et al. 2025). These seven interconvertible species dynamically change in response to extracellular signals (Malek et al. 2017) and cause the recruitment and activation of Rho-type GTPases, actin filament elongating proteins, nucleation promoting factors, as well as severing proteins. Enzymes regulating phosphoinositide metabolism, including PI3K and OCRL, are implicated in autism, epilepsy and intellectual disability, underscoring the potential relevance of these pathways to circuit formation and neurodevelopment (Volpatti et al. 2019).

*In vitro*, phosphoinositide-enriched membranes spontaneously generate filopodia-like actin bundles, highlighting their role as a molecular bridge between acute membrane signalling and actin dynamics, beyond the wider cellular changes caused by phosphoinositide lipid signalling, such as in cell growth, vesicular transport and intracellular calcium release (Lee et al. 2010; Wills & Hammond 2022). In neuronal growth cones, brain-derived neurotrophic factor (BDNF) activates phosphoinositide 3-kinase (PI3K) via TrkB and p75 NT receptors, triggering Cdc42 activation, cofilin dephosphorylation, and the formation of longer, more numerous filopodia (Gehler et al. 2004a,b; Chen et al. 2006; Luikart et al. 2008). Areas of PI(3,4,5)P_3_ enrichment precede F-actin patch formation that leads to protrusion of axonal filopodia (Ketschek & Gallo 2010) and acute production of PI(3,4,5)P_3_ by optogenetic control leads to wave-like growth cone actin morphologies (Kakumoto & Nakata 2013).

Recognised pathways to PI(3,4)P_2_ include class I PI3K production of PI(3,4,5)P_3_ followed by dephosphorylation by 5-phosphatases or via phosphorylation of PI(4)P by class II PI3K (Malek et al. 2017; Zhang et al. 2017). While the enrichment of phosphoinositide lipids at the tips of filopodia suggests they are involved (Jacquemet et al. 2019), how phosphoinositide lipid signalling-based conversions mediate the recruitment and activation of multiple actin regulators that result in specific cytoskeletal architectures and dynamics is not understood.

There are a wide array of candidate proteins that transduce information from phosphoinositide lipids to the actin machinery, including myosin motors, nucleation promoting factors and adaptor proteins. For filopodia, these include lamellipodin, myosin-X, WAVE, TOCA-1, IRSp53 and Cdc42 (Nozumi et al. 2002; Disanza et al. 2006; Lebensohn & Kirschner 2009; Plantard et al. 2010; Umeki et al. 2011; Yoshinaga et al. 2012; Taylor et al. 2019; Cheng & Mullins 2020; Fox et al. 2022; Popović et al. 2022; Pokrant et al. 2023; Blake et al. 2024). Phosphoinositide lipids are recognised to regulate actin remodelling in phagocytosis and they are converted in real-time during the inward curvature of clathrin-coated pits (He et al. 2017; Montaño-Rendón et al. 2022). The development of small molecule inhibitors targeting specific phosphoinositide enzymes (Suwa et al. 2009; Furet et al. 2013; Pirruccello et al. 2014; Wright et al. 2014; Lo et al. 2023), alongside optimised imaging probes that resolve phosphate positions on the inositol ring (Stauffer et al. 1998; Goulden et al. 2019), enables the acute perturbation and visualisation of phosphoinositide conversions at high temporal resolution in diffraction-limited filopodia.

Our current understanding of filopodia is that they integrate multiple molecular pathways, with compositions that are heterogeneous across time and between individual protrusions in a cell (Urbančič et al. 2017; Dobramysl et al. 2021). Based on the constrained and stereotypical patterns of filopodial extension and retraction we observed, our working model is that fluctuating combinations of filopodial proteins provide a robust mechanism for converting extracellular signals into elongated, bundled actin and the slender filopodial morphology. We developed an image analysis pipeline, *Filopodyan*, which allows causal relationships to be drawn between dynamic fluctuations in actin regulators, such as Ena/VASP proteins, and filopodial behaviour, even in the presence of redundancy and heterogeneity (Urbančič et al. 2017). Here, quantitative statistical approaches offer the ability to map the directionality of signalling and effector pathways to filopodial protrusion and retraction and distinguish noise from productive signals (Noh et al. 2022; Blake et al. 2024).

In this work, we combined high-resolution imaging of filopodial dynamics with acute perturbation of phosphoinositide lipid conversions using small molecule inhibitors and lipid-specific probes, to uncover how they influence the dynamic behaviour of axonal growth cone filopodia.

## Results

### Expression of phosphoinositide lipid probes and filopodial metrics

RGC axons emerge from eye primordia dissected at stage 35-36 after overnight culture in minimal conditions when plated onto laminin-coated dishes (Dwivedy et al. 2007). We expressed phosphoinositide lipid binding probes alongside membrane marker GAP43-RFP using RNA injection into 4-cell stage embryos and after explanting eyes, acquired images using an elliptical total internal reflection fluorescent microscope acquiring two wavelengths simultaneously. We used EGFP-PLCδ-PH to detect PI(4,5)P_2_ (Stauffer et al. 1998), TAPP1 EGFP-cPHx3 to detect PI(3,4)P_2_ (Goulden et al. 2019) and tried EGFP PH-BTKx2 with a nuclear export signal (Walpole et al. 2022) and Grp1 PH domain (Kavran et al. 1998) to detect PI(3,4,5)P_3_. PLCδ-PH gives the characteristic plasma membrane staining pattern (Fig. 1A) and TAPP1-3xcPH showed robust localisation to filopodial tips (Fig. 1B). RGCs from PH-BTKx2 and Grp1 PH-injected embryos failed to send out axons and had poor survival, consistent with recent reports that a single molecule approach is needed for PI(3,4,5)P_3_ probes to avoid blocking growth and survival signals (Holmes et al. 2025). RGCs expressing TAPP1-3xcPH had reduced survival and outgrowth compared with PLCδ-PH however at low levels of expression, there were no effects of TAPP1-3xcPH on filopodial dynamics (SI Fig. 1A) while expression of PLCδ-PH led to a modest reduction in filopodial tip retraction (SI Fig. 1B).

**Figure 1.**
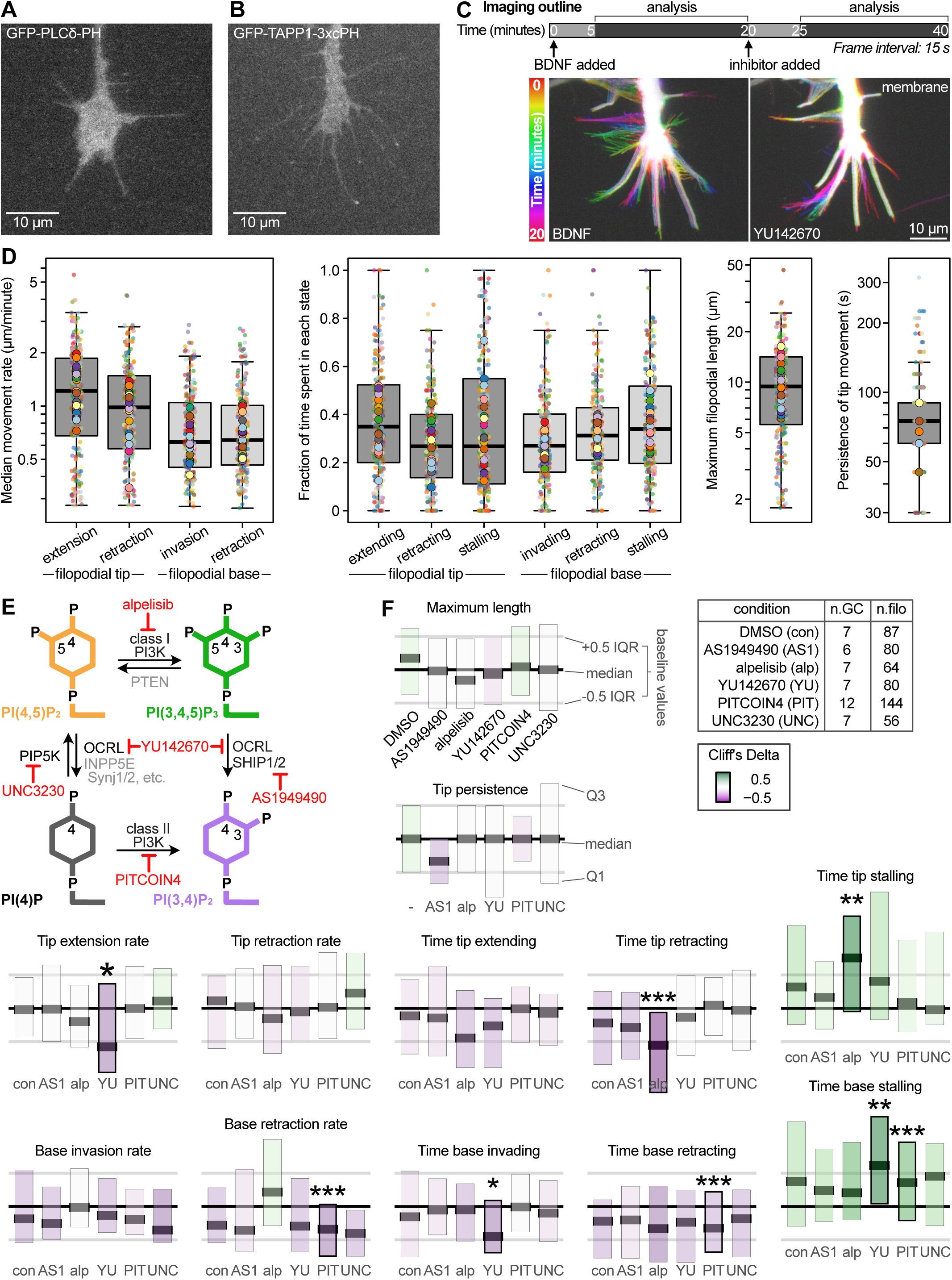
Profiling filopodial dynamics reveals spatially distinct effects of phosphoinositide lipid inhibitors. (A) *Xenopus* Retinal Ganglion Cell (RGC) axonal growth cone expressing GFP-PLCδ-PH showing uniform plasma membrane localisation. (B) RGC expressing GFP-TAPP1-3xcPH showing punctate localisation at filopodial tips. (C) Imaging outline, showing typical 40 minute imaging experiment (see Methods for more details) comprising 20 minutes of BDNF treatment followed by 20 minutes of inhibitor treatment. To allow equilibration, data from minutes 5-20 were analysed. At 15 s frame interval, dynamic filopodia can be quantified under different conditions (time projection shown as rainbow colour scale). (D) Profiling baseline filopodial dynamics parameters shows the range of movement rates of filopodial tips and bases, fraction of time spent moving or stalled, maximum lengths and tip movement persistence (based on the auto-correlation function). (E) Phosphatidylinositol lipid diagram focused on conversions of PI(4,5)P_2_ and PI(3,4,5)P_3_ showing selected kinases and phosphatases and the inhibitors used in this study. (F) Filopodial dynamics parameters after treatment with a panel of inhibitors reveals distinct stalling effects. For each parameter, the boxplot for each inhibitor showing median and lower/upper quartiles was normalised to filopodial dynamics in the same cells before drug addition. The overall central lines show the medians of baseline conditions and the overall grey lines show the interquartile range of baseline conditions. Significant changes from before to after treatment are shown in bold boxes with asterisks denoting p value (* < 0.05, ** < 0.01, *** < 0.001). Significance was assessed by Mann-Whitney tests with multiple comparisons corrected using the Benjamini-Hochberg method for all parameters within each condition. Boxplot colour shows Cliff’s Delta (a measure of effect size) from white (0, no effect) to green/magenta (+1/-1, strong positive/negative effect).

BDNF treatment activates phosphoinositide 3-kinase signalling and is reported to lengthen filopodia via activation of neurotrophin receptors (Gehler et al. 2004a,b). It is also a crucial mediator of branch complexity in response to correlated and non-correlated synaptic vesicle firing in retinotectal axons in tadpoles *in vivo* (Cohen-Cory & Fraser 1994; Kutsarova et al. 2023). Therefore we added BDNF to the culture medium prior to imaging. The frequency of timelapse acquisition was optimised to minimise photobleaching while adequately sampling filopodial dynamics on acute application of the experimental intervention (Fig. 1C). Images were aligned to segment the growth cone shape and filopodia across the time course using the membrane marker, allowing us to quantify multiple aspects of filopodial dynamics.

Growth and retraction rates of the RGC filopodia are approximately exponentially distributed, agreeing with previous observations (Dobramysl et al. 2021), and so we have used a log y axis to present the data in a superplot format (Fig. 1D). Filopodial tip extension and retraction rates range from 0.2-6 µm/min. The rates of movement of filopodial bases are smaller than the tips, agreeing with filopodial tips being the dominant site determining their properties (Mallavarapu & Mitchison 1999). Filopodia spend approximately equivalent periods of time extending, retracting or stalling in the basal conditions, with filopodial lengths and rates determined by both tip and base behaviour (Fig. 1D). Lengths range from 2 µm to more than 25 µm. Filopodial tips typically engage in a pattern of movement for around 75 s (measured by the persistence time, Fig. 1D).

### PI3K inhibitors and OCRL/INPP5B inhibitor YU142670 slow filopodial extension in distinct ways

To elucidate the role of phosphoinositide lipid conversions in filopodial dynamics we used established inhibitors that modulate the key steps between PI(4,5)P_2_, PI(3,4,5)P_3_ and PI(3,4)P_2_ that localise to filopodia (Fig. 1E). We sought to uncouple the effects on transcription and cell growth from dynamic effects on actin cytoskeletal regulators, so used acute treatment. We chose alpelisib as a specific class Iα PI3K inhibitor to target the conversion of PI(4,5)P_2_ to PI(3,4,5)P_3_ (Furet et al. 2013), PITCOIN4 as most recent class IIα PI3K inhibitor developed to inhibit PI(4)P to PI(3,4)P_2_ (Lo et al. 2023), YU142670 that inhibits OCRL and INPP5B to target PI(4,5)P2 to PI(4)P (Pirruccello et al. 2014), UNC3230 to inhibit PI4P 5-kinase PIP5K1C for PI(4)P to PI(4,5)P_2_ (Wright et al. 2014) and AS1949490 that inhibits SHIP2 to target PI(3,4,5)P_3_ to PI(3,4)P_2_ (Suwa et al. 2009).

Baseline datasets were acquired by taking images every 15 s for 15 minutes, then inhibitors were added to the dish and after 5 minutes incubation, imaging was continued to capture the inhibitor effects within the same growth cones. We quantified filopodial parameters before and after DMSO or inhibitor treatment, showing the median and interquartile range of each condition post-treatment normalised to its baseline data, with significance calculated relative to the baseline data (Fig. 1F).

The largest effects on tip dynamics were from alpelisib and YU142670, that decreased the overall movement, exhibiting distinct phenotypic profiles (Fig. 1F). PITCOIN4 and YU142670 significantly reduced base dynamics, both leading to stalling behaviours. This shows that phosphoinositide lipid conversion is important at multiple locations within filopodia that have distinct actin organisations.

The effect of YU142670 increasing stalling at both tips and bases shows that prevention of PI(4,5)P_2_ dephosphorylation reduces actin turnover, consistent with the increased F-actin staining previously observed with YU142670 (Pirruccello et al. 2014). The inhibitors of class I (alpelisib) and class II PI3K (PITCOIN4) both slow or stall filopodial dynamics, acting at different locations (Fig. 1F). RGCs treated with alpelisib, that is expected to decrease conversion of PI(4,5)P_2_ to PI(3,4,5)P_3_ in response to growth factor signalling, showed a significant decrease in the time filopodial tips spent retracting and increased the time they spent stalling (Fig. 1F). When filopodia were extending, the tip extension rate was maintained in the presence of alpelisib (Fig. 1F). This indicates that preventing metabolism of PI(4,5)P_2_ to PI(3,4,5)P_3_ leads to deficiencies in actin turnover, and also that PI(3,4,5)P_3_ may have a role in filopodial retraction. PITCOIN4 reduced the retraction rate and the time the base spent retracting, with an accompanying increase in the time the bases were stalled (Fig. 1E). This suggests there are roles for PI(3,4)P_2_ at the base of filopodia, perhaps in endocytosis or integrin adhesions (Bu et al. 2009; Posor et al. 2013, 2022).

AS1949490 and UNC3230 had no significant effects under these conditions, which could be due to the short incubation times (Fig. 1F). To test whether the BDNF addition was having an acute effect on filopodial dynamics, we used the same imaging protocol, imaging growth cones for 10 minutes prior to BDNF addition, and then monitored the effects between 5 and 20 minutes after BDNF addition on the same growth cones. There were no significant effects on filopodial dynamics (SI Fig. 1C). BDNF application led to an increase in PI(3,4)P_2_ within the whole filopodium and at filopodial tips specifically (SI Fig. 1D). BDNF treatment had no effect on PLCδ-PH fluorescence within filopodia (SI Fig. 1E).

### Non-intuitive effects of alpelisib and YU142670 indicate the complexity of phosphoinositide lipid networks and cytoskeletal regulation

Despite alpelisib and YU142670 both theoretically leading to increases in PI(4,5)P_2_ we have previously found that alpelisib treatment of kidney proximal tubule cells counteracts actin accumulations on endosomes caused by knockout of *OCRL* (Daste et al. 2017; Berquez et al. 2020). This prediction was based on disentangling how phosphoinositide conversion through PI(3,4,5)P_3_ to PI(3)P led to timed co-activation of actin polymerisation through PI(4,5)P_2_ and PI(3)P via adaptor protein SNX9 during endocytosis (Daste et al. 2017). In this model, alpelisib was predicted to decrease actin polymerisation by reducing levels of 3-phosphoinositides. When we stained for phosphoinositide lipids in these experiments, while alpelisib led to an overall drop in PI(3,4,5)P_3_ and PI(3)P, it also decreased staining for PI(4,5)P_2_ (Berquez et al. 2020).

Expression of PLCδ-PH reduced retraction (SI Fig. 1B) suggesting that the probe may compete with PI3K for binding to PI(4,5)P_2_. Despite the expectation that alpelisib and YU142670 treatment should increase plasma membrane PI(4,5)P_2_, we observed no difference in the levels of the PLCδ-PH fluorescence in filopodia on application of either YU142670 or alpelisib (Fig. 2A-B). This could be because these affect local pools or because PI(4,5)P_2_ is present at much higher levels in the plasma membrane than other phosphoinositide lipids, so a relatively small and hard to measure alteration to PI(4,5)P_2_ levels can produce a detectable change in the levels of downstream lipids. PI(4,5)P_2_ levels were also unchanged across the whole growth cone body (Fig. 2C).

**Figure 2.**
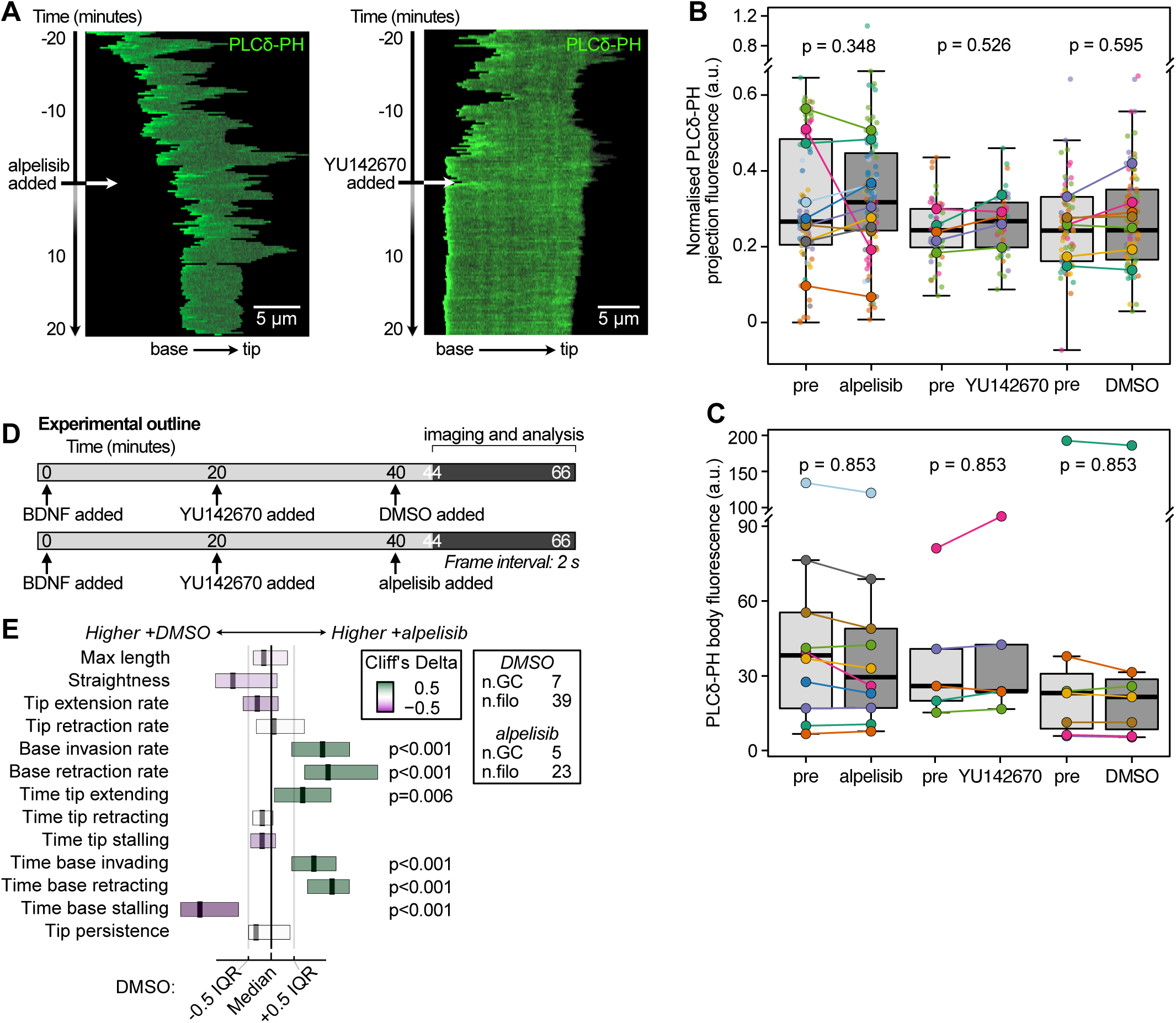
Alpelisib counteracts rather than exacerbates YU142670 effects. (A) Kymograph showing a filopodium (grey outline) as a single pixel line with the tip at the right, time moving downwards, with GFP-PLCο-PH levels shown in green. GFP-PLCο-PH showed an even distribution along the filopodium with some base enrichment. There were no changes in GFP-PLCο-PH levels or distribution after acute alpelisib (left) or YU142670 treatments, while filopodial stalling was apparent. (B) Quantification of mean GFP-PLCο-PH intensity along the filopodium and over time. Boxplots show mean value per filopodium (small dots) and medians per growth cone (large dots), boxes show median and quartiles across all filopodia. Significance assessed by Mann-Whitney tests, with multiple comparisons corrected using the Benjamini-Hochberg method across all dynamics and fluorescence parameters for each inhibitor. (C) Quantification of the mean GFP-PLCο-PH intensity in the growth cone body for each growth cone. Dot colours as in B. (D) Experimental outline showing the combined treatment of RGCs with YU142670 and alpelisib/DMSO. To control for length of time in YU142670, different growth cones were compared between 44-66 minutes, each for 4 minutes measured with 2 s frame interval. (E) Boxplots show dynamics parameters after combined alpelisib treatment, normalised to DMSO treatment for each parameter. Medians and interquartile ranges for DMSO condition shown as central black/grey lines.

Next, we decided to test whether the effects of alpelisib and YU142670 on RGC filopodial dynamics were additive, as would be expected from them both elevating PI(4,5)P_2_, or whether they have opposite effects on filopodia, as expected from our previous work on alpelisib (Daste et al. 2017). In this second scenario, either complex effects on phosphoinositide metabolism are taking place, or the interplay between multiple phosphoinositides results in complex effects on actin regulation. We treated RGCs with YU142670 for 20 minutes, then added either alpelisib or DMSO for 20 minutes, and compared filopodial dynamics (Fig. 2D). While different growth cones are being compared in these experiments to control for the duration of YU142670 incubation, there was no additive effect of alpelisib and YU142670 together, and there was evidence for an alleviation of YU142670-mediated stalling by alpelisib (Fig. 2E).

### Quantitative correlation analysis reveals an association between PI(3,4)P_2_ and persistent filopodial extension

Because the changes in filopodial stalling did not correspond with changes in PI(4,5)P_2_, we took advantage of the ability to analyse native fluctuations in phosphoinositide lipid probes to elucidate the relationship between phosphoinositide conversion and filopodial dynamics.

Manual examination of filopodial tips suggested an association between filopodial protrusion and increased TAPP1-3xcPH signal for PI(3,4)P_2_ (Fig. 3A). Using 2 s timelapse imaging, we performed quantitative correlation analysis between tip extension rate and enrichment of the TAPP1-3xcPH PI(3,4)P_2_ probe at filopodial tips. For each filopodium, we calculated a cross-correlation score between tip fluorescence and tip movement across a range of time offsets (Fig. 3B). This allows for a time delay between filopodial movement and PI(3,4)P_2_ levels in either direction, as would be expected for processes that are indirectly connected. The analysis reveals that almost all filopodia have stronger correlation between TAPP1-3xcPH PI(3,4)P_2_ filopodial tip fluorescence and tip extension rate than expected by chance, and a set of filopodia have strong positive correlation (Fig. 3B-C, SI Fig. 2A-B). The same analysis showed almost no filopodia with significant correlation between PLCδ-PH and tip movement (SI Fig. 2C) or between membrane marker GAP43-RFP signal and tip movement (SI Fig. 2D). TAPP1-3xcPH showed no enrichment over membrane marker during filopodial initiation, despite the slowing in base dynamics induced by class II PI3K inhibitor PITCOIN4 (SI Fig. 2E).

**Figure 3.**
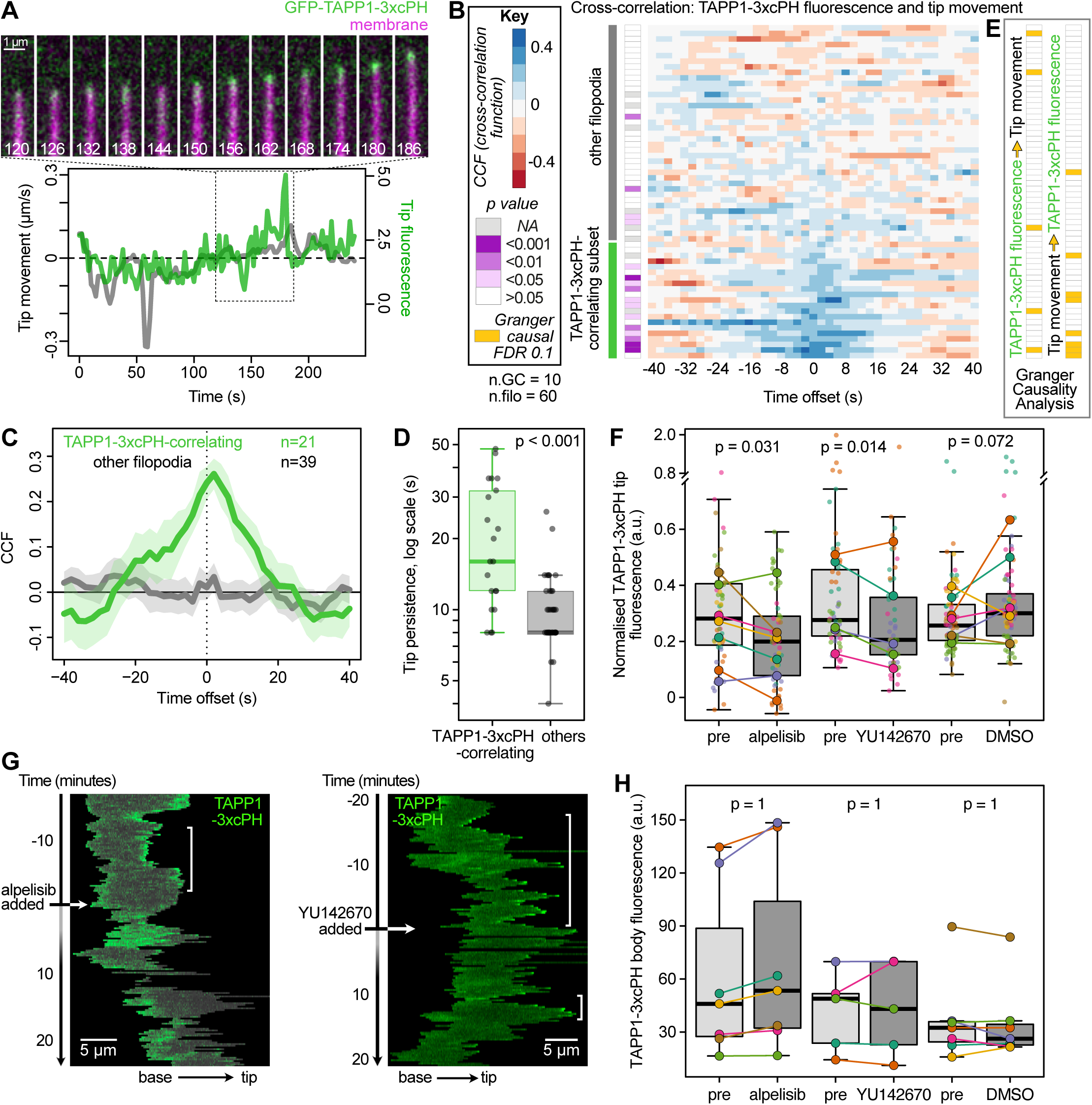
Correlation analysis shows a role for PI(3,4)P_2_ in filopodial protrusion. (A) Montage of an extending filopodium tip, with images acquired every two seconds to capture higher resolution fluctuations in signal, showing correlation between TAPP1-3xcPH signal and filopodium tip extension, taken from a longer time series. Time indicated in seconds. (B) Cross-correlation scores for each filopodium, each calculated at different time offsets, showing a subset with high correlation between TAPP1-3xcPH signal and filopodial tip movement. The heatmap is arranged in order of increasing CCF score averaged over the central seven lags. P values (purple boxes) indicate Markov chain simulation likelihoods of that degree of correlation occurring by chance for each filopodium. (C) Plots showing the average CCF scores for TAPP1-3xcPH-correlating (green) and other filopodia. TAPP1-3xcPH correlation peaks with an offset of 2 s. (D) Boxplot of tip persistence of TAPP1-3xcPH-correlating filopodia compared to the others. Persistence values are lower than baseline dataset in Fig. 1D due to the different frame intervals used. Significance assessed by Mann-Whitney test and corrected for multiple comparisons using the Benjamini-Hochberg method compared to other dynamics parameters. (E) Filopodia with a significant Granger-causal relationship between TAPP1-3xcPH signal and tip movement shown as yellow boxes aligned with heatmap, for each direction of causality. (F) Quantification of mean GFP-TAPP1-3xcPH intensity at the filopodial tips and over time. Boxplots and p values as Fig. 2B. (G) Kymographs showing punctate GFP-TAPP1-3xcPH signal at filopodial tips (regions with white brackets) before alpelisib or YU142670 treatment, and reduced tip localisation after. (H) Quantification of GFP-TAPP1-3xcPH intensity across the growth cone body showing no changes. Boxplots as Fig. 2C.

We defined PI(3,4)P_2_-correlating filopodia as those with a CCF score >75^th^ percentile of simulated data, and these high correlation filopodia had peak correlation with a small positive offset (i.e. the peak TAPP1-3xcPH signal occurs after tip movement; Fig. 3C). These filopodia moved more persistently than other filopodia, while there was no difference in their mean TAPP1-3xcPH levels (Fig. 3D). This suggests that in these more persistent filopodia PI(3,4)P_2_ has a greater relationship with movement: either tip movement is responding to PI(3,4)P_2_ as an upstream signal, or tip movement is driving PI(3,4)P_2_ levels as a downstream response.

Granger causality analysis is a statistical method that can not only establish a causal relationship between two fluctuating processes but also identify the direction of the relationship (Welf & Danuser 2014; Noh et al. 2022). This is particularly powerful in analysing cell biological processes where perturbations can have complex effects on redundant or interconnected processes, such as those we observed with inhibitors (Fig. 1F), where analysing the co-fluctuations in native conditions offers an alternative way to establish cellular mechanisms. To determine Granger causality between tip movement and fluorescence, we compare two autoregression models: tip movement using past fluorescence values and past tip movement values (a full model) with tip movement using past tip movement values only (a reduced model). If the incorporation of fluorescence values improves the predictive power, then fluorescence is Granger causal for tip movement.

Applying Granger causality analysis to this dataset showed that many of the filopodia with strong positive correlation between TAPP1-3xcPH fluorescence and forward movement had significant causality (Fig. 3E, yellow bars compared to dark blue rows on heatmap).

Importantly, contrary to correlation analysis, which produces time lags as a weak indicator of temporal order (because lags are strictly uninformative when the two variables are in a feedback relation), Granger causal inference can identify the directionality of influence.

Exploiting this feature, we observed that in some filopodia, PI(3,4)P_2_ was causal for tip movement (as might be expected from recruitment of lamellipodin being upstream of VASP (Cheng & Mullins 2020)). Unexpectedly, in other filopodia, tip movement was instead causal for increased PI(3,4)P_2_, suggesting that PI(3,4)P_2_ plays multiple roles during filopodial extension at different phases (Fig. 3E).

As PI(3,4)P_2_ has a functional role at filopodial tips, we asked whether the YU142670 and alpelisib treatments (the inhibitors that led to filopodial tip stalling) were affecting the levels of PI(3,4)P_2_. *A priori*, alpelisib might be expected to reduce PI(3,4)P_2_ by reducing levels of PI(3,4,5)P_3_ production. YU142670 could either indirectly increase PI(3,4)P_2_ by providing more PI(4,5)P_2_ substrate or reduce it by inhibiting the dephosphorylation of PI(3,4,5)P_3_ to PI(3,4)P_2_ (Schmid et al. 2004). Acute treatment with either alpelisib or YU142670 reduced levels of PI(3,4)P_2_ at filopodial tips specifically (Fig. 3F-G), while PI(3,4)P_2_ levels across the growth cone were unchanged (Fig. 3H). These experimental perturbations demonstrate a functional association (of either causal direction) between reduced tip extension rates and tip stalling induced by alpelisib and YU142670 treatment and a reduction in PI(3,4)P_2_ levels.

### Bidirectional causality between tip extension and TOCA-1 tip accumulation

To investigate whether a positive feedback loop between filopodial movement and membrane environment is unique to PI(3,4)P_2_ or if it is evident for membrane binding adaptor proteins as well, we attempted to examine the relationship between phosphoinositide lipids and membrane adaptor proteins by analysing lamellipodin, SNX9 and IRSp53. Lamellipodin binds PI(3,4)P_2_ and foci coalesce with VASP to form a complex in the nucleation of B16F1 melanoma cell line filopodia and microspikes (Cheng & Mullins 2020). We were unable to attain expression of fluorescently-labelled lamellipodin in RGCs. While SNX9 shows localisation to some filopodial tips, we did not detect a correlation with filopodial movement (SI Fig. 3A-C). While it could be expressed, the I-BAR and VASP binding protein, IRSp53 (fused with GFP) did not localise to RGC filopodia (SI Fig. 3D). We previously reported a Granger causal relationship between TOCA-1 and filopodial tip protrusion without distinguishing the direction of causality (Blake et al. 2024). We reanalysed the previous TOCA-1 dataset, and show the correlation heatmap ordered by strength of correlation together with the separable directions of Granger causality analysis (Fig. 4A). TOCA-1 correlates with forward movement in about half of cases and like TAPP1-3xcPH, TOCA-1 showed causality in both directions. TOCA-1 accumulation causes tip movement and tip movement causes TOCA-1 accumulation in about 15-20% of filopodia each (Fig. 4A). This shows that multiple, bidirectional molecular mechanisms between membrane environment and filopodial extension are a broad feature of filopodial tips.

**Figure 4.**
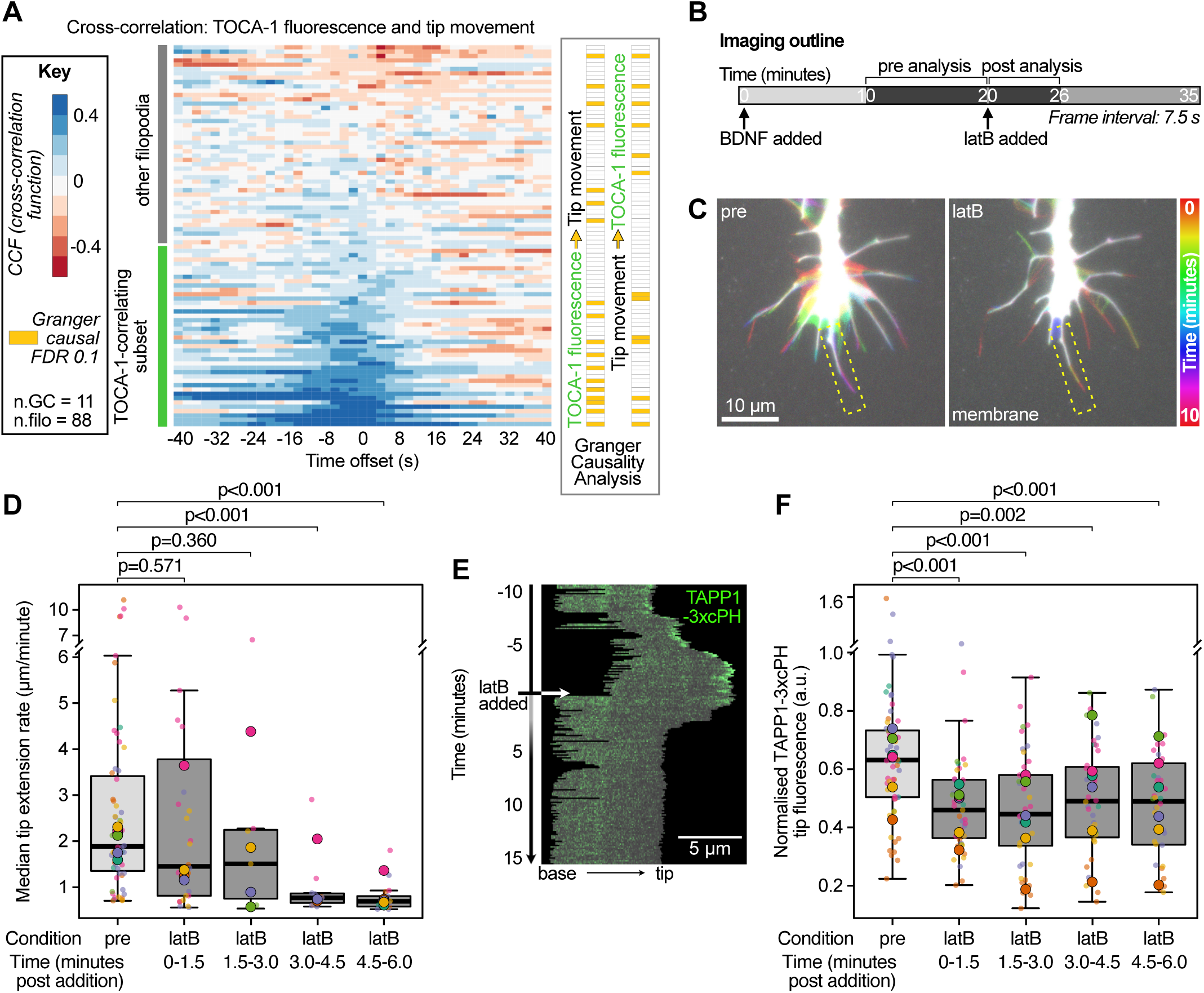
Bidirectional causality is evident for membrane adaptors and depends on actin polymerisation. (A) Cross-correlation heatmap between mNeonGreen-TOCA-1 fluorescence and filopodial tip movement, replotted based on previous data (Blake et al. 2024) with filopodia ordered by CCF scores. Breaking down Granger causality results by direction of causality shows both directions are present at similar rates (yellow boxes), with more overlap between high-correlation filopodia and TOCA-1 causing tip movement. (B) Imaging outline for latrunculin B (latB) experiments. As usual BDNF was added for 20 minutes, then latB was added and imaging continued for 15 minutes, at a 7.5 s time interval as a compromise between duration and rapidity of the response. The drug effect reached full strength after 5-6 minutes. (C) Time projection of a RGC treated before (left) or after treatment with 25 nM latB showing strong filopodial stalling after 10 minutes of treatment. (D) Time course of median tip extension rates before (pre) or after latB treatment at the indicated time windows, showing a significant fall in tip extension rate from 3-6 minutes. Boxplots and p values as in Fig. 2B. (E) Kymograph of GFP-TAPP1-3xcPH fluorescence in a filopodium (yellow dashed boxes in (C)) showing punctate tip localisation before latB addition, with fluorescence is immediately lost after treatment. (F) Time course of mean tip fluorescence per filopodium in the indicated windows, showing a significant drop in the first 90 s of latB treatment.

### Actin polymerisation connects filopodial extension to PI(3,4)P_2_ enrichment

The existing model is that phosphoinositide lipid enrichments act upstream of actin, causing the increased localisation and clustering of membrane adaptors (TOCA-1, lamellipodin, IRSp53 etc.) that in turn interact and cluster the actin filament elongators and bundlers (Disanza et al. 2006; Yoshinaga et al. 2012; Taylor et al. 2019; Blake et al. 2024). In that case, we would have expected the PI(3,4)P_2_ increase induced by BDNF (SI Fig. 1E) to cause persistent filopodial movement, which it did not, suggesting either that there are functionally independent PI(3,4)P_2_ pools at filopodial tips or that BDNF has wider effects that suppress the contributions of PI(3,4)P_2_. Moreover, our observations with TAPP1-3xcPH and TOCA-1 fluorescence with movement indicate that forward movement induces accumulation of membrane adaptors and PI(3,4)P_2_. To test this, we monitored the PI(3,4)P_2_ in the presence of latrunculin B (latB), to prevent actin monomer incorporation at the tips of filopodia. If PI(3,4)P_2_ is upstream of actin polymerisation and filopodial movement then we would expect TAPP1-3xcPH fluorescence to remain stable at filopodial tips, whereas if tip movement or actin polymerisation causes PI(3,4)P_2_ production or inhibits its conversion to other lipids, TAPP1-3xcPH fluorescence should be reduced by latB treatment. RGCs were treated with a low dose (25 nM) of latB, which led to filopodial tip extension gradually stalling over 5 minutes (Fig. 4B-D). By contrast, TAPP1-3xcPH was rapidly lost from filopodial tips within the first 90 s (Fig. 4E-F). This was specific to the tips, as TAPP1-3xcPH fluorescence was not altered across either the growth cone body (SI Fig. 3E) or the filopodium as a whole (SI Fig. 3F). TAPP1-3xcPH fluorescence was lost prior to filopodial stalling, indicating that new actin incorporation directly stimulates PI(3,4)P_2_ synthesis at filopodial tips (Fig. 4E-F).

## Discussion

By combining acute treatment with phosphoinositide lipid enzyme inhibitors, live imaging of fluorescent phosphoinositide reporters and quantitative filopodial dynamics analysis, we have demonstrated that PI(3,4)P_2_ plays multiple roles upstream and downstream of tip protrusion at axonal growth cone filopodia. Perturbing phosphoinositide lipid signalling with PI3K or OCRL inhibition using alpelisib and YU142670 impaired filopodial protrusion and PI(3,4)P_2_ levels. Though OCRL is typically implicated in removing the 5’ phosphate of PI(4,5)P_2_, the ability of OCRL inhibition to reduce PI(3,4)P_2_ levels here could result from OCRL dephosphorylating PI(3,4,5)P_3_ as seen *in vitro* though not previously reported *in vivo* (Schmid et al. 2004). The phosphoinositide enzyme perturbations we used showed some counter-intuitive behaviours. Applying alpelisib after YU142670 did not exacerbate the filopodial tip stalling phenotype and instead there was some improvement. It may be that the inhibitors also led to changes in PI(4,5)P_2_ levels that were not detectable with the EGFP-PLCδ-PH probe, causing reduced actin incorporation which in turn could lead to the decrease in PI(3,4)P_2_.

This agrees with Granger causality analysis suggesting a bidirectional causal relationship between PI(3,4)P_2_ and tip extension and the decrease in PI(3,4)P_2_ observed with latB treatment. The class II PI3K inhibitor PITCOIN4 and YU142670 slowed filopodial base dynamics during the course of the filopodial lifetime, although we did not see an accumulation of TAPP1-3xcPH at the sites of new filopodial initiation.

At filopodial tips, our observations indicate multifaceted mechanisms in which, firstly, upstream PI(3,4)P_2_ production promotes actin polymerisation, perhaps via lamellipodin, which has been shown to regulate VASP activity at the tip of microspike actin bundles (Yoshinaga et al. 2012; Cheng & Mullins 2020). Secondly, filopodial protrusion can induce further production of PI(3,4)P_2_, potentially leading to an increase in persistence of filopodial movement, consistent with the two-fold increase seen in TAPP1-3xcPH-correlating filopodia. Our results also do not preclude PI(3,4,5)P_3_ or PI(3)P being the active species.

The molecular nature of this actin-based or mechanosensitive link from filopodial protrusion to phosphoinositide metabolism is an area of future study. The link appears to be restricted to filopodial tips as latB treatment did not affect PI(3,4)P_2_ levels elsewhere. The growing actin bundle or force applied to the plasma membrane could activate phosphoinositide lipid enzymes directly or indirectly, such as by curvature sensitive phosphatidylinositol kinases and phosphatases or small GTPase activation. Complex non-linear effects within phosphoinositide cascades are increasingly being recognised and our work adds the intersection with actin dynamics to the possible mechanisms (Fung et al. 2024).

Since most, but not all, of the filopodia that showed significant Granger causality only demonstrated one or the other directions of causality, this could reflect filopodia with different phases of activity that demonstrate the respective causal relationships. Our data show consistent correlation between PI(3,4)P_2_ and tip movement on short time scales (Fig. 3B and SI Fig. 2A), and imply that there is a shift in molecular mechanism at longer time scales where PI(3,4)P_2_ can either be upstream or downstream of actin. LatB treatment affected PI(3,4)P_2_ levels within 90 s, giving an upper bound on the timescale of mechanism switching. Longer duration imaging followed by Granger causality analysis will help resolve whether these two mechanisms are distinct or form a feedback loop.

The downstream signalling from filopodial tip protrusion to PI(3,4)P_2_ could also serve another purpose than directly stimulating the growth of the actin bundle, such as regulating filopodial point contact adhesions (Woo et al. 2009; Posor et al. 2022) or vesicular trafficking events, which in turn could support persistent filopodial growth (He et al. 2017; Nozumi et al. 2017; Li et al. 2021).

In conclusion, we show that multiple interventions cause changes in PI(3,4)P_2_ at axonal growth cone filopodial tips, implicating BDNF signalling, class I PI3K and OCRL in its control. PI(3,4)P_2_ at filopodial tips both causes filopodial protrusion and results from it, leading to persistent extension behaviour. Whether these mechanisms constitute a feedback loop and how they relate to adhesion and calcium signalling in filopodia is an important future question (Gomez et al. 2001; Woo et al. 2009). These insights establish a mechanistic basis for filopodial persistence and define spatially distinct mechanisms of actin regulation that inform models of neuronal miswiring and circuit assembly.

### Materials and methods Plasmids and RNA

TAPP1-3xcPH (NES-EGFP-cPHx3) was amplified using Phusion High-Fidelity polymerase (NEB) from NES-EGFP-cPHx3, a gift from Gerry Hammond (Addgene plasmid #116855; http://n2t.net/addgene:116855; RRID:Addgene_116855) using primers containing an FseI site 5’-GCATGGCCGGCCACCATGGCTCTGCAGAAAAAGTTGG-3’ and an AscI site 5’-GGCGCGCCCCGCGGTACCGTCGACTCAGC-3’. Following digestion, the PCR product was cloned into FseI and AscI cut pCS2 his FA plasmid, constructed as previously described (Urbančič et al. 2017). PLCδ-PH domain was amplified from pCDNA 3.1 EGFP-PLCδ-PH which was a gift from Seth Field, Harrington Discovery Institute, Cleveland, Ohio with primers 5’-GATCGGCCGGCCGCTATCGATACCATGGACTCGGGCCGG-3’ and 5’-

GGCGCGCCTCACTCCTTCAGGAAGTTC-3’ which have FseI and AscI sites respectively and cloned into FseI/AscI cut pCS2 GFP FA. NES-EGFP-PH-BTKx2 was amplified from NES-EGFP-PH-BTKx2, a gift from Sergio Grinstein (Addgene plasmid #183652; http://n2t.net/addgene:183652; RRID:Addgene_183652). Primers with FseI and AscI sites 5’-GCATGGCCGGCCGCCACCATGGCTCTGCAGAAAAAGTTGG-3’ and 5’-GGCGCGCCGATCAGTTATCTAGATCCGGTGGATCC-3’ were used and the digested PCR product was cloned into FseI and AscI cut pCS2 his FA. GAP43-RFP was described previously (Urbančič et al. 2017). Capped RNA was transcribed from NotI digested plasmids using SP6 mMESSAGE mMACHINE (Invitrogen) and purified by RNeasy Kit (Qiagen). *X. laevis* SNX9 was PCR amplified from pCMV-sport6-SNX9 (IMAGE clone 3402622; Source Bioscience) using primers with a FseI 5’-GCATGGCCGGCCACCATGAACAGCTTTGCGG-3’ and AscI site 5’-GGCGCGCCTCACATCACTGGG-3’. The digested PCR product was cloned into FseI and AscI cut pCS2 his mNeonGreen FA plasmid, previously described in (Urbančič et al. 2017).

*X. laevis* Brain-Specific Angiogenesis Inhibitor 1-Associated Protein 2-Like 1 (IRSp53) was PCR amplified from pCMV-sport6-IRSp53 (IMAGE clone 7981694, Open Biosystems) using primers 5’ -GCATGGCCGGCCACCATGTCCCGGGACGCAG-3’ and 5’-GGCGCGCCTCATCGGATGATTGGCG-3’ which have FseI and AscI sites respectively. The digested PCR product was cloned into FseI and AscI cut pCS2 his mNeonGreen FA plasmid.

### Xenopus RGC explants

Xenopus embryos were obtained by in vitro fertilisation. 4 cell embryos were micro-injected with 75 pg probe RNA and 75 pg GAP-RFP RNA into the two blastomeres with neural fates of 4 cell embryos. Embryos were cultured in 0.1x Modified Marc’s Ringers (MMR) and staged according to Nieuwkoop and Faber. RGC explants were taken at stage 35 – 36 and cultured overnight in 60% Leibovitz’s L-15 Medium with 100 U/ml penicillin, 100 µg/ml streptomycin and 250 µg/ml amphotericin at room temperature on 35 mm glass bottom dishes (MatTek P35G-1.5-14-C) which had been coated with 10 µg/ml poly-L-lysine (Merck) for 1 hour and 10 µg/ml laminin (Merck) for 10 minutes.

### Live imaging of RGC growth cones

Images were obtained with HILO illumination using a custom built TIRF microscope based on the Nikon Ti Eclipse equipped with an iLas2 illuminator (Roper Scientific) and either a Multisplit beam splitter (Cairn Research) and a Kinetix CMOS camera or with an Optosplit beam splitter (Cairn Research) and a CMOS camera (Hamamatsu ORCA-Flash4.0). Images were typically acquired at 15 s frame interval for a total of 40 minutes. BDNF was added to a concentration of 50 ng/ml and individual RGC growth cones were imaged for 20 minutes, then inhibitors were added and RGCs were imaged for 20 further minutes. Inhibitors were added to final concentrations of: 50 µM YU142670 (Merck), 10 µM alpelisib (BYL719) (Selleck Chemicals), 10 µM AS1949490 (TOCRIS), 20 µM PITCOIN4 (MedChemExpress LLC) or 50 µM UNC3230 (TOCRIS). Data from minutes 5-20 of BDNF treatment were compared to data from minutes 5-20 of inhibitor treatment. For quantifying the effect of BDNF, RGCs were imaged for an additional 10 minutes before BDNF addition, and data from this period was compared to 5-20 minutes after BDNF addition. For the combined treatment of YU142670 and alpelisib, BDNF and YU142670 were added for 20 min each, then either alpelisib or DMSO was added, with images acquired at a 2 s frame interval for 4 minutes during the window of 4-26 minutes after alpelisib or DMSO addition with different growth cones imaged in each condition. For correlation and causation analysis between PI(3,4)P_2_ and dynamics, the frame intervals were 2 s to best calculate the relationships between dynamics and fluorescence. For latB experiments, RGCs were treated with BDNF for 20 minutes, followed by latB treatment for 15 minutes with a frame interval of 7.5 s.

Images were compared from the same RGCs for the final 10 minutes of BDNF treatment to latB treatment at indicated time periods.

### Image analysis

Image processing and analysis was carried out as previously (Blake et al. 2024). Videos were screened for inclusion in analysis based on several criteria: a minimum intensity of protein of interest (15 grey values above background), growth cones that were not static or unhealthy before imaging in baseline conditions, growth cones that were not visibly impacted by contact with another cell, or debris or other issues preventing accurate segmentation. This resulted in typical data acquisitions of 3-5 growth cones per eye, 1-2 of eyes per session leading to analysis of 5-10 movies per condition. Typically the levels of GAP43-RFP and phosphoinositide probe had comparable intensities and no bleedthrough. For fluorescence measurements, values were subject to a background correction based on signal near the growth cone boundary and normalisation to the growth cone body were performed (except for growth cone body fluorescence values, which were just background corrected). This controls for variable expression levels and photobleaching. In general, normalisation reduced the magnitude of changes after drug treatment rather than creating changes. Filopodial tip and base coordinates were smoothed with a moving average of window 5 (for 15 s frame interval measurements) or 3 (for 2-7.5 s frame interval measurements) and removal of outliers (top and bottom 0.5% of values). For quantification of filopodial dynamics and fluorescence parameters, time series with at least 12 frames were used, except for the latB-treated cells where the threshold was set at 4 frames due to the short time windows used in the time course. Correction for multiple comparisons was carried out using the Benjamini-Hochberg method per dataset (i.e. across all dynamics and fluorescence parameters measured for one inhibitor treatment), except for mean body fluorescence which is a per-growth cone metric rather than per-filopodium. There, the correction was applied for each set of p values appearing on the same graph. Cross-correlation analysis was calculated for all filopodia with at least 50 frames, without removal of missing values.

### Granger causality analysis

Conceptually, to assess the Granger causality from time series X to time series Y, we compare two autoregressive models: a full model and a reduced model. In the full model, values of Y are predicted using past values of X and past values of Y. In the reduced model values of Y are predicted using past values of Y only. If the full model provides a significant improvement in predictive power compared to the reduced model, then X is said to Granger cause Y. Granger causality analysis was conducted similarly to our previous analysis (Blake et al. 2024). We confirmed the stationarity of the time series by using the Augmented Dickey Fuller Test (Dickey & Fuller 1979), then for the Granger causality test we selected the optimum lag from 1 to 5 that minimised the Bayesian Information Criteria in the full model (Schwarz 1978). We used the MATLAB function gctest() to carry out the Granger causality test, and corrected raw p values for multiple hypothesis testing using the Benjamini-Hochberg false discovery rate procedure (Benjamini & Hochberg 1995).

## Acknowledgements

We would like to thank Matthias Krause, Gerry Hammond and Sergio Grinstein for the kind gift of their plasmids, Richard Butler for discussions about Filopodyan and Raghu Padinjat for providing feedback on the manuscript. This work was funded by a Wellcome Trust Senior Research Fellowship 219482/Z/19/Z to JLG.

## Supplementary Information Figure legends

**SI Figure 1. Dynamics and fluorescence effects of phosphoinositide lipid probe expression and inhibitors.** (A) Expression of TAPP1-3xcPH does not affect filopodial dynamics, compared to GAP43-RFP membrane marker alone. RGCs imaged every 2 s for 4 minutes. (B) Expression of GFP-PLCδ-PH reduced filopodial retraction compared to the same GAP43-RFP data, while other dynamics parameters were unchanged. (C) Comparing RGCs before and after BDNF treatment showed no significant dynamics changes. Significance tests and normalised boxplots as in Fig. 1. Significance shown when p<0.05. (D) BDNF treatment significantly increased mean GFP-TAPP1-3xcPH fluorescence at filopodial tips and across the filopodium. (E) BDNF treatment did not alter GFP-PLCδ-PH intensity across the filopodium.

**SI Figure 2. Additional correlation analysis for TAPP1-3xcPH, PLCδ-PH and membrane marker levels during filopodial initiation.** (A) TAPP1-3xcPH and tip movement CCF scores are significantly higher than random, based on a bootstrap approach - block reshuffling the time series data for tip movement and recalculating cross-correlation with tip fluorescence to create a simulated dataset. The actual correlation scores (averaged from -6 s to +6 s time offsets) at a range of indicated quantiles is shown as red bars, while the distribution of 1000 simulated datasets is shown as violins. Mann-Whitney tests were used to assess if the actual data was significantly different, and the Benjamini-Hochberg method was used to correct for multiple comparisons (* < 0.05, ** < 0.01, *** < 0.001). Here, the actual correlation was significantly stronger than random for all quantiles. (B) For comparison with PLCδ-PH and GAP43-RFP membrane marker correlation results, the TAPP1-3xcPH correlation analysis was repeated after removing all datapoints where filopodium length <1.5 µm, because in those other datasets, correlation scores were unreliable when length <1.5 µm. The degree of TAPP1-3xcPH correlation was robust to this additional filter. (C) Cross-correlation heatmap and comparison to simulated data for PLCδ-PH, showing almost no correlating filopodia. (D) Similar results for GAP43-RFP membrane marker. (E) For filopodia expressing TAPP1-3xcPH, where possible the future filopodial base (defined as the closest point on the cell surface to the future site of initiation) was tracked before filopodial initiation for up to 40 s. GFP-TAPP1-3xcPH and membrane marker intensities were plotted separately, showing that TAPP1-3xcPH signal did not deviate from the membrane marker pattern before initiation. Lines show means, shaded areas show 95% CI.

**SI Figure 3. Investigating putative membrane binding adaptor proteins and TAPP1-3xcPH signal during latB treatment.** (A) SNX9 shows diffuse/weakly punctate localisation in RGCs and filopodia (white arrows). (B) Filopodia were imaged at 7.5 s frame interval, and data were processed with a moving average filter (window size 5). Time series with more than 17 frames were used for cross-correlation analysis. (C) A bootstrap reshuffling approach showed no significant correlation with filopodial tip movement. (D) IRSp53 shows only diffuse localisation in RGCs, without clear membrane localisation, preventing quantitative analysis. Images representative of 11 growth cones. (E) After latB treatment, mean GFP-TAPP1-3xcPH fluorescence in the growth cone body did not change. Boxplots as in Fig. 2C. (F) After latB treatment, mean GFP-TAPP1-3xcPH fluorescence across the whole filopodium does not change.

**Video S1. TAPP1-3xcPH is dynamically localised to filopodial tips**. RGC growth cone showing dynamic filopodia with TAPP1-3xcPH (green) at tips, especially during extension. Plasma membrane labelled by GAP43-RFP membrane marker (magenta). 4 minute video acquired at 2 s frame interval, replayed at 10 frames per second.

**Video S2. TAPP1-3xcPH is rapidly lost from filopodial tips after low dose latrunculin B treatment.** RGC growth cone showing dynamic filopodia with TAPP1-3xcPH (green) at tips. Once latB is added (t = 0), TAPP1-3xcPH is lost from filopodial tips within 90 s, while filopodial stalling increases over a longer time period.

