## Supplementary figures and images for "Phosphoinositide lipids have bidirectional and spatially distinct roles in filopodial dynamics"

### SI Fig. 1

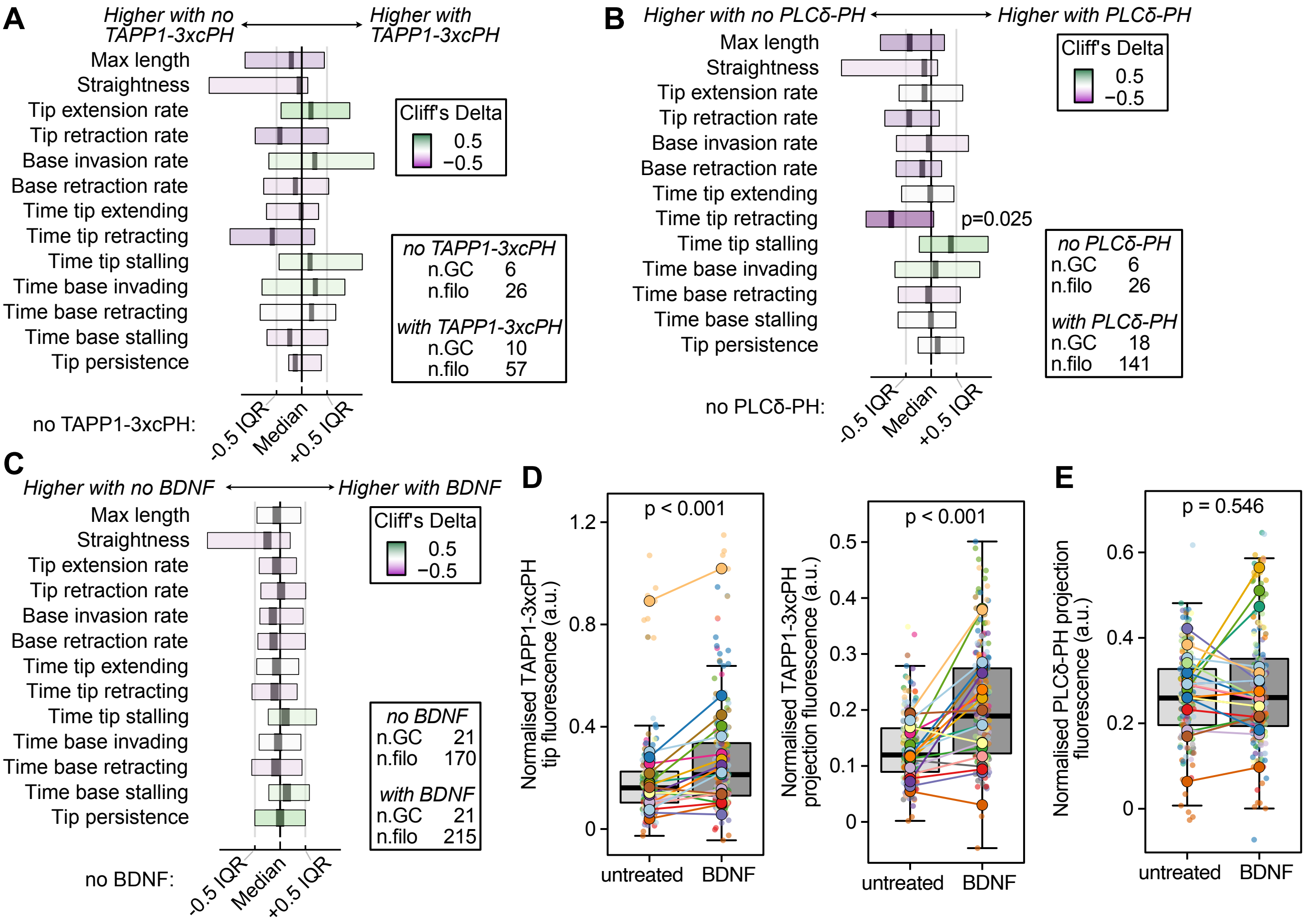

### SI Fig. 2

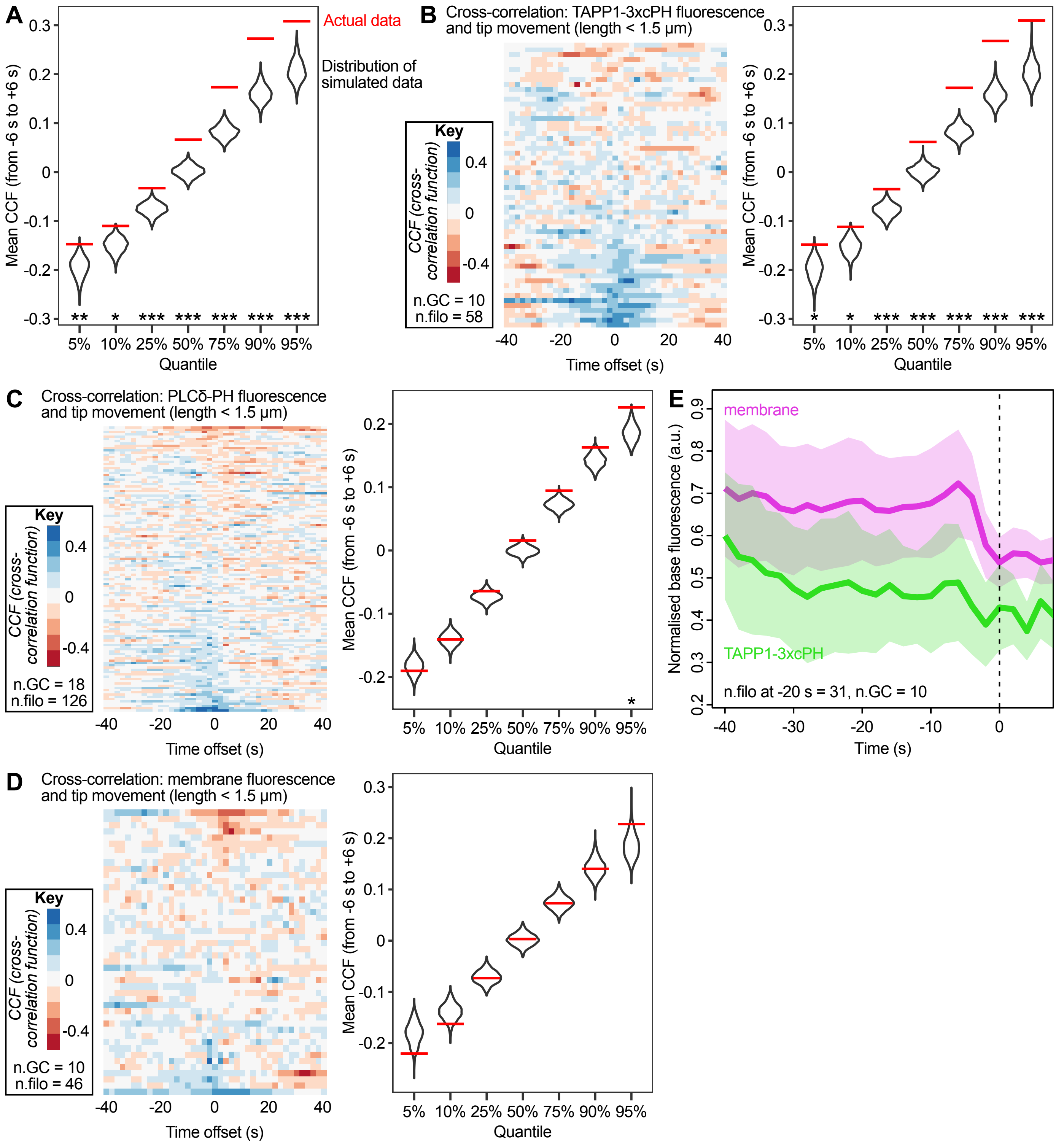

### SI Fig. 3

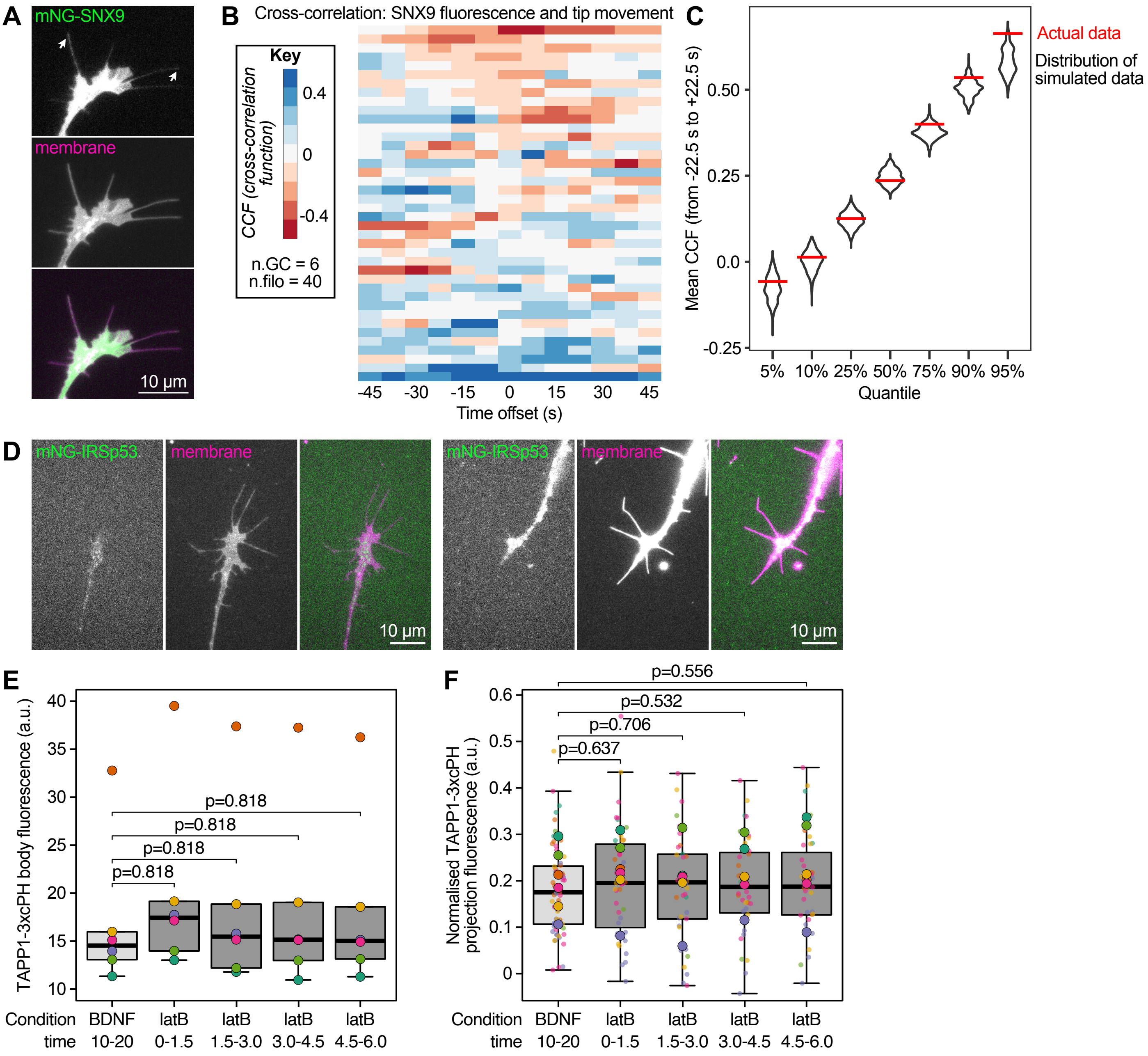
